# Molecular interplay between peptidoglycan integrity and outer membrane asymmetry in maintaining cell envelope homeostasis

**DOI:** 10.1101/2025.06.19.660527

**Authors:** Sinjini Nandy, Arshya F. Tehrani, Augusto C. Hunt-Serracin, Jacob Biboy, Christine Pybus, Waldemar Vollmer, Joseph M. Boll

## Abstract

The bacterial cell envelope is a critical interface with the environment, particularly in Gram-negative species where the outer membrane and peptidoglycan layers coordinate to maintain structural integrity and resist turgor. Although this coordination is essential for survival, the molecular mechanisms linking outer membrane and peptidoglycan homeostasis remain poorly understood. LD-transpeptidases (LDTs) are enzymes that crosslink peptides in peptidoglycan and incorporate D-amino acids, but their physiological roles are not fully defined. Here, we characterize the activity of the LDT enzyme LdtJ in *Acinetobacter baumannii* and investigate the consequences of its deletion. Loss of LdtJ disrupts cell morphology, downregulates peptidoglycan precursor genes (e.g., *dadA*, *alr*), and activates the stringent response, including elevated ppGpp levels and *dksA* upregulation. These defects are fully suppressed in a Δ*ldtJ* Δ*mla* double mutant, implicating the outer membrane lipid transport Mla pathway in compensatory regulation. RNA sequencing revealed that transcriptional changes in the Δ*ldtJ* mutant are reversed in the double mutant, highlighting a functional interplay between peptidoglycan remodeling and outer membrane lipid asymmetry. Our findings suggest that LdtJ contributes to envelope integrity not only through peptidoglycan modification but also by influencing broader regulatory and metabolic networks.

**IMPORTANCE:** *Acinetobacter baumannii* is a leading cause of hospital-acquired infections and is highly resistant to antibiotics. Its survival relies on the integrity of the cell envelope, composed of the peptidoglycan layer and outer membrane. While LD-transpeptidases (LDTs) are traditionally known for reinforcing peptidoglycan structure through non-canonical crosslinking, our findings reveal that the LdtJ enzyme also plays a critical role in regulating cellular metabolism and stress responses. Deletion of *ldtJ* results in pronounced growth defects and abnormal cell morphology – phenotypes that are fully suppressed by disrupting the outer membrane lipid asymmetry transport system, Mla. This genetic interaction uncovers a previously unrecognized link between peptidoglycan remodeling and outer membrane lipid homeostasis. These insights deepen our understanding of envelope coordination in Gram-negative bacteria and suggest that targeting interconnected stress response pathways could offer novel strategies to undermine bacterial resilience.

## INTRODUCTION

The Gram-negative bacterial cell envelope is composed of three distinct layers: the inner (cytoplasmic) membrane, the peptidoglycan (PG) layer located in the periplasm, and the outer membrane (OM). The PG is a mesh-like polymer that maintains cell shape, resists osmotic pressure, and protects against environmental stressors [1,2]. Coordination between the PG and OM is essential for preserving envelope integrity and bacterial viability. Understanding how these layers interact and respond to stress is key to elucidating mechanisms of bacterial survival and morphology. PG is composed of glycan chains made of repeating N-acetylglucosamine (Glc*N*Ac) - N-acetylmuramic acid (Mur*N*Ac) disaccharides, connected by short peptides. In many Gram-negative bacteria, these peptides typically includes L-alanine, D-glutamate, *meso*-diaminopimelic acid (*m*DAP), and two D-alanine residues, with the peptide covalently linked to Mur*N*Ac [3–5]. PG biosynthesis and remodeling requires a suite of enzymes, including glycosyltransferases and transpeptidases for PG polymerization and cross-linkage, and PG hydrolases – such as endopeptidases, carboxypeptidases, amidases, lytic transglycosylases, and lysozymes – for maturation, turnover and degradation [6–9].

Among enzymes involved in PG biosynthesis, penicillin binding proteins (PBPs) play a crucial role. Class A and B PBPs possess DD-transpeptidase activity (DD-TPase), which forms 4-3 peptide crosslinks. These 4-3 crosslinks, formed from the fourth D-alanine of the donor pentapeptide moiety to the third *m*DAP of the acceptor stem [6,10], are the most abundant, comprising 60-100% of total PG crosslinks during growth (depending on species) [1,3]. In contrast, LD-transpeptidases (LDTs) generate LD-crosslinks by linking the third *m*DAP residues of adjacent peptide stems [11–13]. Unlike PBPs, LDTs utilize tetrapeptides as donor substrates, which are generated by DD-carboxypeptidases (DD-CPases) that remove the terminal D-alanine from pentapeptides. While 4-3 crosslinks dominate during exponential growth in *Escherichia coli*, LD-crosslinks typically comprising ∼10% of total crosslinks during exponential growth increase to 16% in stationary phase [14] and can reach up to 30-40% under certain OM stress conditions [15].

Although LDTs are not essential for survival of most bacteria, they perform a variety of functions beyond PG crosslinking [12]. These include D-amino acid (DAA) incorporation into PG [16], maintenance of cell envelope integrity during lipopolysaccharide (LPS) transport defects [15], β-lactam resistance [17,18], cold shock [19] and autolysin regulation [7]. In *A. baumannii*, LdtJ is the only periplasmic LDT, suggesting it catalyzes LD-crosslink formation and DAA incorporation, particularly during the stationary phase. A previous study [20] showed that deletion of LdtJ leads to reduced growth and altered cell morphology. While wild-type cells exhibited a coccobacillary shape, the Δ*ldtJ* mutant appeared coccoid and significantly a shorter cell length, a phenotype that could be fully complemented. Although LDTs are typically associated with stationary-phase functions, our prior work suggested that LdtJ also contributes to growth-phase physiology. However, the molecular mechanism underlying these roles remain unclear and merit further investigation.

The Gram-negative OM is an asymmetric barrier, with phospholipids enriched in the periplasmic inner leaflet and LPS/lipooligosaccharide (LOS) present in the surface-exposed outer leaflet. This lipid asymmetry is critical for the cell to maintain the OM barrier function that protects against many external agents such as detergents, antibiotics or lysins, which can disrupt membrane integrity or cause cell lysis [21,22]. Under stress conditions, phospholipids can become mislocalized to the outer leaflet, compromising the asymmetry. To counteract this, Gram-negative bacteria employ mechanisms to restore lipid distribution and preserve OM integrity [21,23]. In *A. baumannii*, two conserved systems help to maintain OM asymmetry: the Mla (maintenance of OM lipid asymmetry) pathway and phospholipase A (PldA) [24]. The Mla system consists of six proteins – MlaA, MlaB, MlaC, MlaD, MlaE, and MlaF – that work together to remove mislocalized phospholipids from the OM and transport them back to the inner membrane [21,24]. In *E. coli*, MlaA, an OM lipoprotein, interacts with OmpC to capture mislocalized phospholipids [25]. MlaC then shuttles them to the inner membrane MlaFEDB complex, where MlaF provides ATPase-driven energy for transport [26–29]. In the absence of a functional Mla pathway, phospholipids accumulate on the cell surface, disrupting membrane asymmetry and compromising barrier integrity. As a result, *mla* mutants exhibit hypersensitivity to membrane-disrupting agents such as SDS and EDTA, which destabilize the OM by chelating divalent cations and disrupting lipid packing [21,24].

The PG has long been considered the primary structural component of the bacterial cell envelope, providing mechanical strength and maintaining cell shape [5,30]. However, recent studies have shown that the LPS-enriched OM also plays a critical role in shaping cell morphology and reinforcing envelope integrity, particularly in resisting internal turgor changes [31–34]. Despite these advances, the molecular mechanisms that coordinate the functions of the PG layer and OM to maintain homeostasis remain poorly understood.

Herein, we characterized the enzymatic functions of LdtJ in *A. baumannii*. We confirmed its LD-transpeptidase and DAA incorporation activities and further identified LD-carboxypeptidase activity. Using the catalytically inactive mutant LdtJ_C390S_, we demonstrated that these enzymatic functions are not required for maintaining growth fitness or cellular morphology under standard conditions. While LdtJ was only thought to mediate LD-crosslinks and DAA incorporation during stationary phase, our findings suggest it also actively contributes to PG synthesis and cellular fitness during active growth. Beyond PG structural remodeling, LdtJ may influence envelope-associated stress responses and metabolic pathways, pointing to a broader regulatory role in maintaining cellular homeostasis – identified for the first time in *A. baumannii*. Notably, we found that introducing the *ldtJ* deletion into a strain lacking the Mla pathway – thereby disrupting OM lipid asymmetry – restored growth-phase defects, including impaired viability and altered morphology. These results underscore the importance of coordinated regulation between PG biosynthesis and OM homeostasis. Together, our findings reveal a functional interplay between PG integrity and OM lipid asymmetry, revealing how crosstalk between these systems supports envelope stability in *A. baumannii*.

## RESULTS

### Activities of the LD-transpeptidase LdtJ

Previous research on LdtJ showed that the gene product contributed to *A. baumannii* fitness and growth [20]. The Δ*ldtJ* mutant cells were spherical, unable to incorporate the fluorescent DAA, and had a growth defect relative to the wild-type strain. To expand on the previous report, we purified the wild-type enzyme and the catalytically inactive version (LdtJ_C390S_), in which the active-site cysteine was substituted with serine. Purification yielded two soluble protein products, as shown with Coomassie staining (**Figure S1A**). Western blots with α-LdtJ specific antisera [20] and an α-His antibody also showed there were two LdtJ protein products (**Figure S1B and S1C**). MS analysis of the two excised bands showed the molecular weights (MW) were 45.59 kDa and 43.38 kDa (**Figure S1D**). The MWs were consistent with the full-length LdtJ_His8X_ protein and a protein with a cleaved signal sequence between A21 and A22, consistent with di-alanine motif cleavage [35,36]. In general, cleavage of the di-alanine-containing signal sequence is associated with successful export of proteins to the periplasm. However, the presence of both cleaved and uncleaved forms suggests that LdtJ is inefficiently exported in *E. coli* strain BL21.

*In vitro* enzymatic assays with recombinant wild-type and active-site mutant enzymes were conducted using PG isolated from an *E. coli* BW25113Δ6LDT strain devoid of all LDT proteins (**Figure 1**), as previously described [33]. Wild-type LdtJ exhibited three distinct activities: LD-transpeptidase, DAA addition, and low-level LD-carboxypeptidase activity (**Figure 1A**). In the absence of LdtJ, Tetra, TetraTetra, and TetraTetraTetra muropeptides were the main muropeptides released from the PG of *E. coli* BW25113Δ6LDT, consistent with previous PG analysis [18]. LdtJ without addition of D-lysine produced distinct LD-crosslinked TriTri(Dap), TetraTri(Dap), TetraTriTri(Dap), and TetraTetraTri(Dap), indicating LdtJ-mediated LD-LD-crosslinking. Additionally, we detected the muropeptides Tri and TetraTri, consistent with low-level LD-carboxypeptidase activity that removes the terminal D-alanine from tetrapeptides to generate tripeptides. Notably, the detection of TriTri(Dap) further supports dual LD-carboxypeptidase and LD-transpeptidase activities of LdtJ, as the formation of this species requires both the formation of LD-crosslinks forming TetraTri(Dap) and the trimming of its tetrapeptide to the tripeptide. Upon the addition of D-lysine, new peaks corresponding to Tetra-D-Lys and TetraTetra-D-Lys (D-Lys replacing D-Ala at position 4) were observed, indicating that LdtJ can incorporate DAAs into PG. However, in the presence of NaCl, the abundance of these D-Lys-containing muropeptides was significantly reduced compared to conditions without salt, suggesting that elevated ionic strength interferes with LdtJ-mediated DAA incorporation. Salt is known to modulate protein conformation and activity by altering electrostatic interactions [37]. Under high-salt conditions, only the LD-crosslinked TetraTri(Dap) and TetraTetraTri(Dap) were detected, further indicating that D-Lys incorporation is selectively impaired. In contrast, LdtJ_C390S_ exhibited no detectable activity on PG (**Figure 1B**). Collectively, these findings indicate that LdtJ possesses both LD-transpeptidase activity and low-level LD-carboxypeptidase activity, along with the ability to mediate D-lysine (DAA) incorporation into PG.

**Figure 1.**
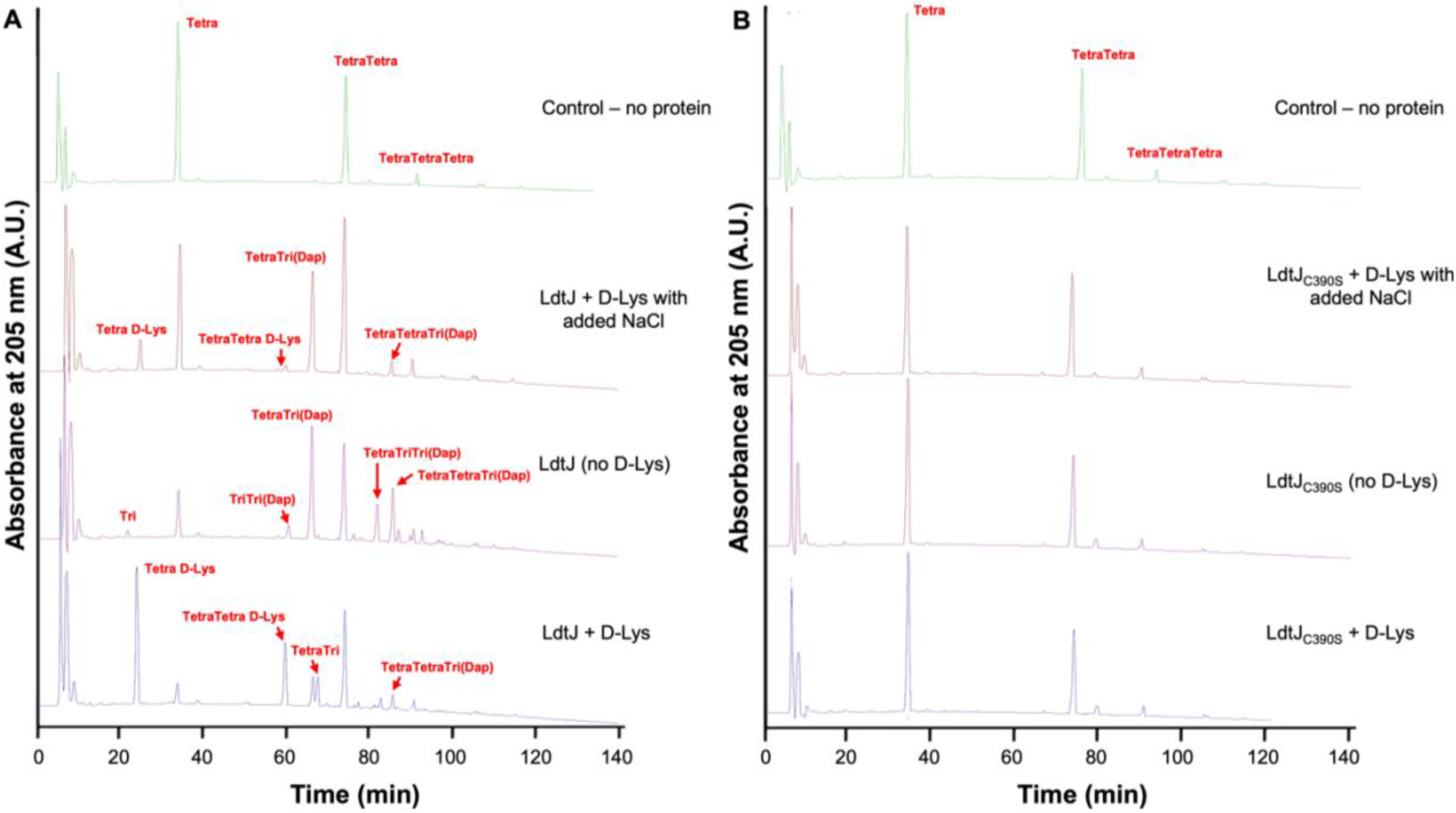
High-performance liquid chromatography (HPLC) traces of LdtJ enzymatic activity. **(A)** HPLC profiles of recombinant LdtJ incubated with purified peptidoglycan (PG) from *E. coli* BW25113Δ6LDT. Digestion with the muramidase cellosyl yields Tetra, TetraTetra, and TetraTetraTetra as the major monomeric, dimeric and trimeric muropeptides (disaccharide peptide PG fragments). Reactions were performed under three conditions: with excess D-lysine (D-Lys), without D-Lys, and with D-Lys plus NaCl. **(B)** HPLC traces of the catalytically inactive mutant LdtJ_C390S_ under the same conditions. Muropeptide peaks are labeled accordingly. The observed profiles are consistent with LdtJ-mediated incorporation of D-amino acids (Tetra-D-Lys, TetraTetra-D-Lys), formation of LD-crosslinks (TriTri(Dap), TetraTri(Dap), TetraTriTri(Dap), TetraTetraTri(Dap)), and low-level LD-carboxypeptidase activity (Tri, TriTri(Dap), TetraTri). The absence of these modifications in the LdtJ_C390S_ mutant confirms the requirement of the catalytic cysteine for enzymatic function.

### Loss of LdtJ enzymatic activity has no impact on fitness and morphology of *A. baumannii* during the growth phase

The LD-transpeptidase, LdtJ, encodes four distinct domains following its signal peptide (**Figure 2A**): 1) an N-terminal region (residues 1-146), 2) a PG binding domain (147-192), 3) an internal linker region (193-282), and 4) the YkuD LDT domain (283-410), which includes the active site cysteine at position 390. To investigate the contribution of each domain in LdtJ stability, we generated a series of complementation constructs expressing LdtJ variants lacking individual domain. Western blot analysis (**Figure 2B**) showed that deletion of the N-terminal region (LdtJ_Δ40-146_), PG binding domain (LdtJ_Δ147-192_), and the internal region (LdtJ_Δ193-282_), resulted in unstable proteins. In contrast, YkuD domain deletion (LdtJ_Δ283-390_/LdtJ_ΔYkuD_), despite removing the catalytic site, produced a stable protein, albeit at reduced levels relative to wild-type LdtJ. However, expression levels were like those observed for wild-type LdtJ and the catalytically inactive mutant LdtJ_C390S_ used in complementation assays.

**Figure 2.**
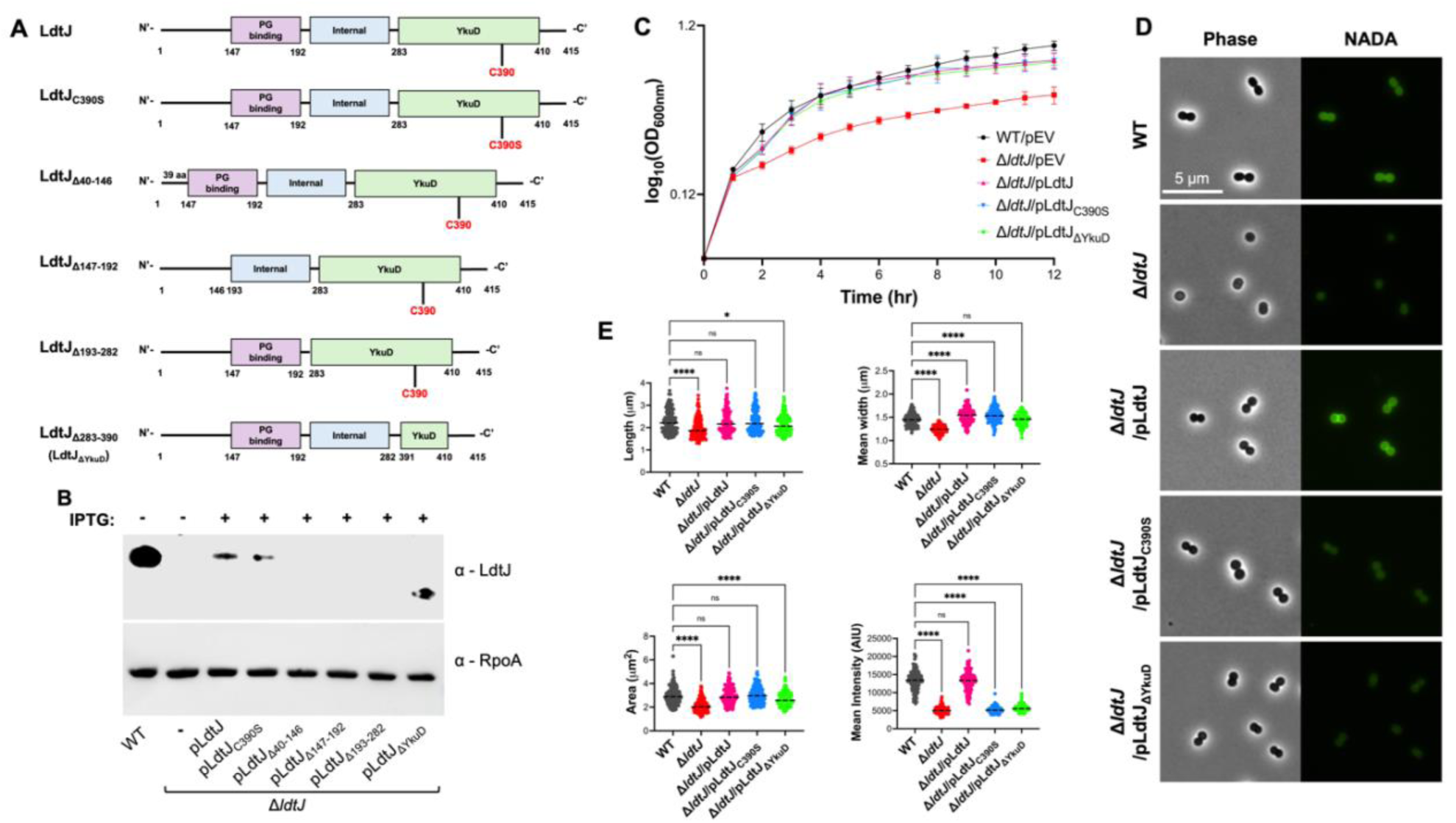
Disruption of enzymatic activity of LdtJ does not impair growth fitness or cell morphology. **(A)** Schematic representation of wild-type (WT) LdtJ and mutant constructs used in this study. **(B)** Western blot analysis using α-LdtJ antibody showing expression of LdtJ (44.5 kDa), LdtJ_C390S_ (44.49 kDa), and LdtJ_ΔYkuD_ (33.07 kDa). α-RpoA was used as a loading control. **(C)** Growth curve analysis of WT, Δ*ldtJ*, and complementation strains over 12 hours in a 24-well plate (*n* = 3) (EV = Empty vector). **(D)** Phase-contrast and NADA-stained microscopy images of WT, Δ*ldtJ*, and complementation strains. **(E)** Quantification of cell dimensions (length, width, and surface area) and intensity for WT, Δ*ldtJ*, and complementation strains (150 < *n* < 200 cells per strain). Each experiment was independently replicated three times, and one representative data is shown. Statistical significance was determined using one-way ANOVA (*p* < 0.05).

Based on a previously reported growth defect in the Δ*ldtJ* mutant [20], we hypothesized that the active site mutant LdtJ_C390S_ and LdtJ_ΔYkuD_ would impair bacterial fitness during growth. Consistent with prior findings, our analysis revealed a significant fitness defect in Δ*ldtJ* compared to the wild-type (**Figure 2C**), which was fully rescued by complementation restored to wild-type levels LdtJ allele. Surprisingly, complementation with either LdtJ_C390S_ or LdtJ_ΔYkuD_ also restored fitness to wild-type levels, suggesting that the LD-transpeptidase and LD-carboxypeptidase activity of LdtJ are not required for maintaining bacterial fitness under these conditions.

In addition to fitness defects, the Δ*ldtJ* strain exhibited altered cell morphology (**Figure 2D**). Consistent with previous findings [20], wild-type cells displayed a coccobacillus shape, whereas Δ*ldtJ* appeared more coccoid. The morphological defect was rescued by complementation with the wild-type *ldtJ* allele, as well as with the LdtJ_C390S_ and LdtJ_ΔYkuD_ variants. Quantitative analysis confirmed that Δ*ldtJ* cells had significantly reduced cell dimensions, including shorter length, narrower width, and smaller surface area compared to the wild-type cells (**Figure 2E**). The wild-type phenotype was fully restored upon complementation with all three alleles. As expected, fluorescence intensity of NADA (a fluorescent DAA) was not restored in strains complemented with LdtJ_C390S_ and LdtJ_ΔYkuD_.

These results indicate that LdtJ is essential for maintaining both bacterial fitness and proper cell morphology. Although the YkuD domain, which contains the active site, is required for LdtJ enzymatic activities, its inactivation does not compromise protein stability, bacterial fitness, or cell shape. This suggests that LdtJ contributes to these cellular functions through a mechanism independent of its catalytic activity.

### Disruption of the maintenance of lipid asymmetry (Mla) pathway compensates for the growth and morphological defects observed in the Δ*ldtJ* mutant

Given the severe growth defect in the Δ*ldtJ* mutant during exponential growth phase, we sought to identify genetic suppressors that could restore this phenotype. To this end, we analyzed a previously published transposon mutagenesis dataset comparing Δ*ldtJ* and wild-type strains [20,38]. This analysis revealed an enrichment of transposon insertions in all genes of the Mla pathway, with four genes showing statistically significant increases in insertion frequency in the Δ*ldtJ* background (**Figure 3A**). We further compared specific transposon insertions in *mlaA::Tn* and *mlaE::Tn* in wild-type and Δ*ldtJ* strains (**Figure 3B**). These findings led us to hypothesize that disruption of the Mla pathway suppresses the growth defect associated with the Δ*ldtJ* mutant.

**Figure 3.**
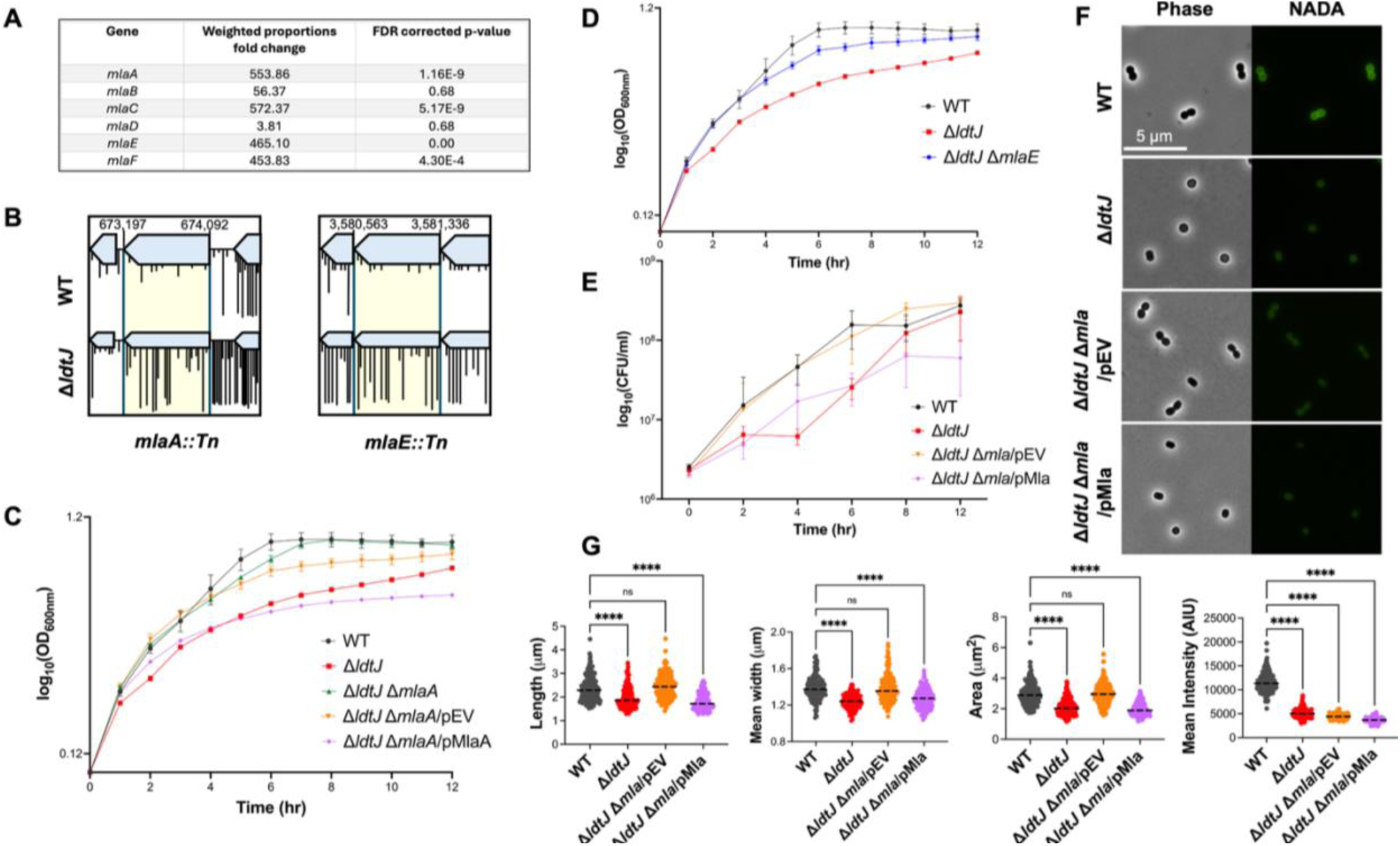
Disruption of the Mla pathway restores growth fitness, viability, and morphology in the Δ*ldtJ* mutant. **(A)** Transposon sequencing analysis of WT and Δ*ldtJ* strains. **(B)** Transposon insertion profiles for *mlaA::Tn* and *mlaE::Tn* in WT and Δ*ldtJ* backgrounds. **(C)** Growth curves of WT, Δ*ldtJ*, and Δ*ldtJ* Δ*mlaA* double mutants, with and without an empty vector, and with Mla pathway complementation. **(D)** Growth curves of WT, Δ*ldtJ*, and Δ*ldtJ* Δ*mlaE* strains. **(E)** Colony-forming unit (CFU) counts for WT, Δ*ldtJ*, Δ*ldtJ* Δ*mla* double mutants with empty vector, and with Mla complementation. **(F)** Phase-contrast and NADA-stained microscopy images of WT, Δ*ldtJ*, Δ*ldtJ* Δ*mlaA* double mutants with empty vector, and with Mla complementation. **(G)** Quantification of cell dimensions and fluorescence intensity from panel F (*n* = 150–200 cells). All experiments were independently replicated three times; one representative data is shown. Statistical significance was determined using one-way ANOVA (*p* < 0.05).

To test this hypothesis, we constructed a panel of double mutants in the Δ*mla* background. Both Δ*ldtJ* Δ*mlaA* and Δ*ldtJ* Δ*mlaE* strains restored growth fitness comparable to wild-type levels (**Figure 3C-3D**). Furthermore, *mlaA* complementation in the Δ*ldtJ* Δ*mlaA* mutant (**Figure 3C**) reverted the growth phenotype back to that of the Δ*ldtJ* single mutant, confirming that the suppression of the growth defect is dependent on disruption of the Mla pathway.

Interestingly, SDS/EDTA susceptibility assays (**Figure S2A**) revealed variability between independent Δ*ldtJ* Δ*mlaA* replicates. One replicate, derived from an SDS/EDTA-resistant Δ*mlaA1* parent, retained resistance, while the other, derived from an SDS/EDTA-sensitive Δ*mlaA2* parent, remained susceptible. Despite these differences in outer membrane susceptibility, both Δ*ldtJ* Δ*mlaA* strains successfully restored the growth defect observed in the Δ*ldtJ* mutant (**Figure 3C, S2B**). Moreover, complementation of *mlaA* in the SDS/EDTA-sensitive Δ*ldtJ* Δ*mlaA2* strain reverted the phenotype to that of the Δ*ldtJ* single mutant (**Figure S2B**). These findings indicate that restoration of Δ*ldtJ* fitness is independent of OM integrity, and instead likely results from disruption of lipid asymmetry. For all subsequent experiments, the SDS/EDTA-resistant Δ*ldtJ* Δ*mlaA1* strain – hereafter referred to as Δ*ldtJ* Δ*mla* – was used as the representative double mutant.

In addition to the Mla pathway, transposon mutagenesis also revealed a high frequency of transposon insertions in *pldA* (**Figure S2C**), a gene encoding a phospholipase involved in maintaining OM lipid asymmetry. To determine whether *pldA* disruption could similarly suppress the Δ*ldtJ* growth defect, we generated the Δ*ldtJ* Δ*pldA* double mutant in a Δ*pldA* background. This strain fully restored growth fitness to wild-type levels (**Figure S2D**). Moreover, the Δ*ldtJ* Δ*mla* Δ*pldA* triple mutant also rescued the Δ*ldtJ* growth defect, further supporting a link between OM asymmetry and suppression of the Δ*ldtJ* phenotype.

Beyond restoring growth, inactivation of Mla in the Δ*ldtJ* Δ*mla* mutant also corrected defects in cell viability (**Figure 3E**) and morphology (**Figure 3F**). The double mutant exhibited wild-type cell length, mean width, and surface area, fully rescuing the spherical morphology observed in Δ*ldtJ* cells (**Figure 3G**). Together, these findings support the conclusion that disruption of OM lipid asymmetry – rather than general defects in membrane integrity, compensates for the fitness, viability, and morphological defects caused by loss of the LD-transpeptidase LdtJ.

### Transcriptomic profiling reveals compensatory gene expression changes in Δ*ldtJ* and Δ*ldtJ* Δ*mla* mutants during growth

To investigate how disruption of the Mla pathway restores growth fitness, viability, and morphology in the Δ*ldtJ* mutant, we performed transcriptomic profiling during the logarithmic growth phase. RNA was isolated from wild-type, **Δ***ldtJ,* and **Δ***ldtJ* **Δ***mla* strains, and gene expression profiles were compared. Differential expression analysis was conducted using a fold-change threshold of ≥ 2 for upregulation or ≤ −2 for downregulation, with two-tailed *t*-test (*p*-value < 0.05). Volcano plots comparing wild-type vs. **Δ***ldtJ* and **Δ***ldtJ* vs. **Δ***ldtJ* **Δ***mla* revealed similar patterns of differential gene expression between these two comparisons. In contrast, the wild-type vs. **Δ***ldtJ* **Δ***mla* comparison showed fewer genes meeting the differential expression criteria, suggesting that Mla disruption partially restores the transcriptomic profile toward a wild-type state (**Figure 4A, 4G-4H**).

**Figure 4.**
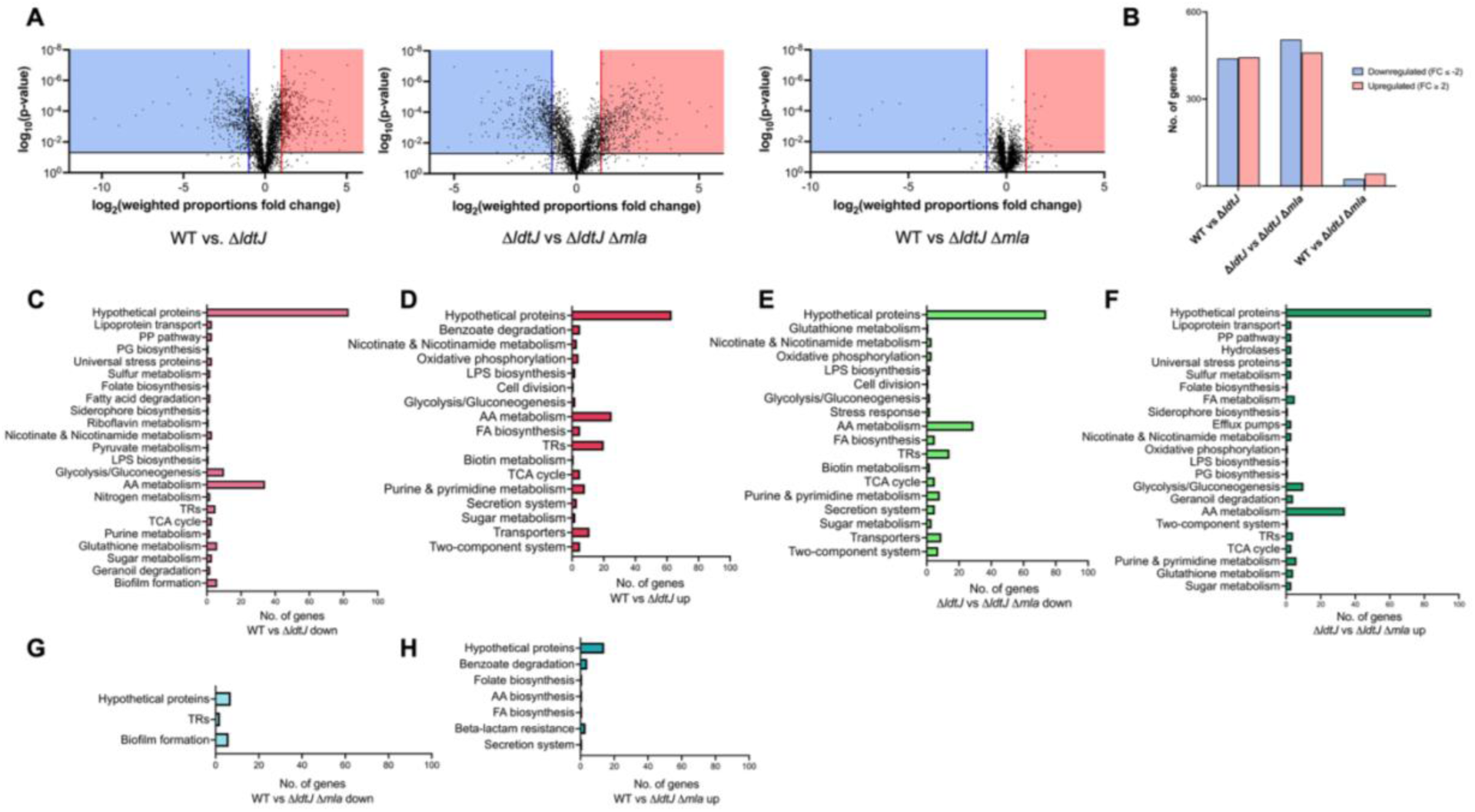
Differential gene expression profiles in WT, Δ*ldtJ*, and Δ*ldtJ* Δ*mla* strains. **(A)** Volcano plot showing differentially expressed genes during the growth phase. The blue box highlights significantly downregulated genes, while the red box highlights significantly upregulated genes. **(B)** Total number of significantly regulated genes, defined by fold change ≥ 2 or ≤ −2 and *p* < 0.05. **(C–H)** Number of differentially expressed genes involved in various biological pathways (e.g., cell envelope biogenesis, stress response) across the following comparisons: WT vs. Δ*ldtJ*, Δ*ldtJ* vs. Δ*ldtJ* Δ*mla*, and WT vs. Δ*ldtJ* Δ*mla* (PP = Pentose phosphate, PG = Peptidoglycan, LPS = Lipopolysaccharide, AA = Amino acid, TR = Transcriptional regulator, TCA = Tricarboxylic acid, FA = Fatty acid).

Compared to the wild-type, the Δ*ldtJ* mutant exhibited substantial transcriptomic changes, with 440 genes downregulated and 444 upregulated (**Figure 4B**). In Δ*ldtJ* vs. Δ*ldtJ* Δ*mla,* 505 genes were downregulated and 460 were upregulated, indicating a similarly broad shift in gene expression. In contrast, the Δ*ldtJ* Δ*mla* strain showed only 25 genes downregulated, and 43 upregulated relative to the wild-type. These results suggest that the transcriptomic profile of Δ*ldtJ* Δ*mla* closely resembles that of wild-type, the conclusion from our phenotypic analyses that Mla disruption compensates for the loss of LdtJ.

Notably, 363 of the 440 genes downregulated in Δ*ldtJ* relative to wild-type were upregulated in Δ*ldtJ* Δ*mla* compared to Δ*ldtJ* (**Figure 4C-F**). These genes were enriched in pathways related to amino acid metabolism, fatty acid degradation, and biofilm formation. Conversely, 387 of the 444 genes upregulated in Δ*ldtJ* were downregulated in Δ*ldtJ* Δ*mla*, including genes involved in nutrient transport and metabolism, fatty acid biosynthesis, stress response, fructose metabolism, and the bacterial secretion system. These reciprocal expression patterns suggest that Mla disruption rebalances key metabolic and stress response pathways disrupted by the loss of LdtJ.

The inverse gene expression patterns observed between wild-type vs. **Δ***ldtJ* and **Δ***ldtJ* vs. **Δ***ldtJ* **Δ***mla* suggest the involvement of a global regulatory mechanism. These findings indicate that the loss of *ldtJ* activates compensatory transcriptional programs, which are further modulated by disruption of the Mla pathway. This layered regulatory response likely underlies the restoration of fitness, viability, and morphology in the **Δ***ldtJ* **Δ***mla* mutant.

### Metabolic and regulatory disruptions in Δ*ldtJ* are mitigated by loss of Mla function during exponential growth

To validate the RNA-sequencing (RNA-seq) results, we performed reverse transcriptase relative fold PCR (RT-relative fold PCR) on DNase-treated RNA samples from wild-type, Δ*ldtJ*, and Δ*ldtJ* Δ*mla* strains. Complementary DNA (cDNA) was synthesized and amplified using gene-specific primers targeting transcripts identified as differentially expressed in the RNA-seq dataset. We selected *dadA*, *alr*, and *argO* for validation based on their significant expression changes and functional relevance – *dadA* and *alr* are associated with D-alanine metabolism and PG synthesis [39,40], while *argO* encodes an arginine exporter involved in maintaining intracellular arginine homeostasis and modulating cellular stress responses [41]. In the Δ*ldtJ* mutant, *dadA* (*A1S_0095*) and *alr* (*A1S_0096*) were significantly downregulated relative to wild-type, with fold changes of 41.0 and 26.9, respectively (**Table S1**). Conversely, *argO* (*A1S_1046*) was markedly upregulated by 32.9-fold. Interestingly, in the Δ*ldtJ* vs. Δ*ldtJ* Δ*mla* comparison (**Table S2**), an inverse expression pattern was observed: *dadA* and *alr* were upregulated by 45.1 and 39.6-fold, respectively, while *argO* expression was reduced by 31.9-fold compared to the Δ*ldtJ* mutant. These relative fold PCR results corroborated the RNA-seq data, confirming the significant downregulation of *dadA* and *alr*, and upregulation of *argO* in the Δ*ldtJ* mutant compared to wild-type (**Figure 5A**). In the wild-type vs. Δ*ldtJ* Δ*mla* comparison (**Table S3**), fold changes for *dadA*, *alr*, and *argO* were below the differential expression threshold indicating expression levels comparable to those in the wild-type background. These findings were consistent with RT-relative fold PCR results, further supporting the conclusion that Mla disruption restores gene expression toward the wild-type state.

**Figure 5.**
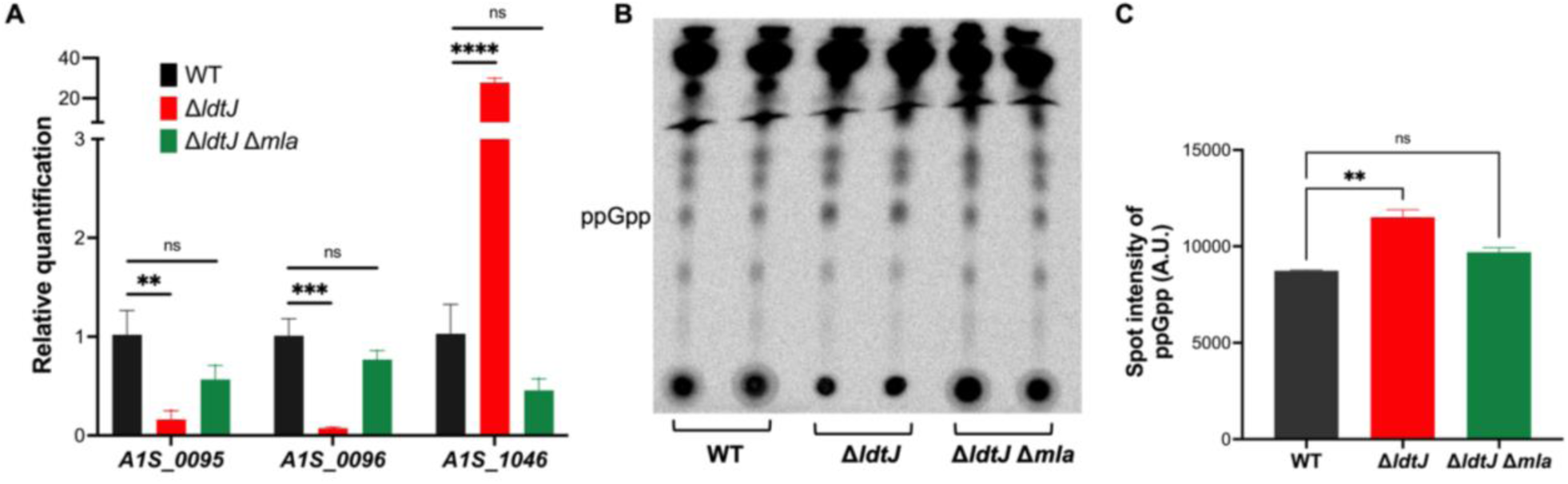
Reverse Transcriptase (RT)-relative fold PCR analysis and ^32^P-radiolabeled thin-layer chromatography (TLC). **(A)** RT-relative fold PCR analysis showing relative expression levels of *A1S_0095*, *A1S_0096*, and *A1S_1046* in WT, Δ*ldtJ*, and Δ*ldtJ* Δ*mla* strains. RT-relative fold PCR experiments were independently replicated three times; one representative dataset is shown. Statistical significance was determined using a two-tailed *t*-test (*p* < 0.05). **(B)** TLC of radiolabeled (^32^P) lipid extracts from WT, Δ*ldtJ*, and Δ*ldtJ* Δ*mla* strains. ppGpp was identified based on migration relative to known standards using a 1.5 M KH₂PO₄ (pH 3.4) solvent system. **(C)** Quantification of TLC spot intensities. TLC experiments were independently replicated twice in duplicate; one representative dataset is shown. Error bars represent standard deviations. Statistical significance was determined using one-way ANOVA (*p* < 0.05).

In addition to transcriptional validation, phenotypic confirmation of the RNA-seq results was obtained by examining the expression of *dksA* (*A1S_0248*), a transcriptional regulator involved in the bacterial stringent response. *dksA* expression was found to be upregulated by 2.8-fold in the Δ*ldtJ* mutant and downregulated by 3.1-fold in Δ*ldtJ* Δ*mla*. *dksA* modulates RNA polymerase activity and functions in concert with the alarmone (p)ppGpp – primarily guanosine-5′, 3′-tetraphosphate, ppGpp – to regulate the transcription of stress response genes, including those involved in PG remodeling under nutrient-limiting conditions [42]. While DksA does not directly bind ppGpp, it is essential for mediating its effect on transcriptional reprogramming.

To investigate the functional link between *dksA* expression and the stringent response, we quantified intracellular (p)ppGpp levels using radiolabeled phosphate (^32^P) followed by thin layer chromatography (TLC). TLC analysis revealed elevated accumulation of ppGpp in the Δ*ldtJ* mutant compared to both wild-type and Δ*ldtJ* Δ*mla* (**Figure 5B**). Quantification of TLC spot intensities (**Figure 5C**) confirmed this observation, indicating an enhanced stringent response in the Δ*ldtJ* background.

In summary, both RT-relative fold PCR and phenotypic validation confirm that deletion of *ldtJ* disrupts the expression of genes associated with PG synthesis and activates stress response pathways. This is evidenced by altered expression of *dadA*, *alr*, *argO*, and *dksA*, along with elevated intracellular ppGpp levels. These disruptions reflect a broader imbalance in PG-related and unrelated metabolic and regulatory networks. Strikingly, additional deletion of *mla* in the Δ*ldtJ* Δ*mla* mutant restores transcriptional and physiological homeostasis, mitigating the stress phenotypes observed in the Δ*ldtJ* single mutant. Together, these findings reveal a functional interplay between OM lipid homeostasis and PG remodeling in *A. baumannii* and suggest that perturbations in one system can be compensated by adaptive changes in the other.

## DISCUSSION

In this study, we uncover a previously unrecognized functional interplay between peptidoglycan (PG) remodeling and outer membrane (OM) lipid homeostasis in *Acinetobacter baumannii*, centered on the LD-transpeptidase LdtJ. Building on prior work linking *ldtJ* to cell shape and growth [20], we demonstrate that LdtJ contributes to both enzymatic and non-enzymatic functions essential for maintaining envelope integrity. Deletion of *ldtJ* resulted in severe growth and morphological defects, disrupted D-alanine metabolism, and activated stress responses, including elevated ppGpp levels. Remarkably, disruption of the Mla lipid transport system fully suppressed these phenotypes, revealing a compensatory relationship between PG remodeling and OM lipid asymmetry.

Biochemical assays confirmed that LdtJ catalyzes LD-PG crosslinking via meso-diaminopimelic acid (*m*DAP) and incorporates D-amino acids (DAAs) such as D-lysine, consistent with the activity of LDTs in *Escherichia coli* and *Mycobacterium tuberculosis* [43,44]. LdtJ also exhibited low-level LD-carboxypeptidase activity, and the catalytic cysteine was essential for function, as shown by the loss-of-function phenotype in the LdtJ_C390S_ variant [45].

Our mutagenesis and complementation analyses revealed that non-catalytic domains of LdtJ are critical for protein stability or folding, while deletion of the catalytic YkuD domain preserved growth and morphology. These findings suggest that LdtJ has non-enzymatic roles, potentially serving as a structural or scaffolding protein. This dual-function model is supported by similar observations in *Francisella tularensis*, *Campylobacter jejuni*, and *Mycobacterium smegmatis*, where PG-modifying enzymes influence cell shape independently of catalytic activity [46–48].

To understand how Mla disruption suppresses Δ*ldtJ* defects, we constructed double and triple mutants. Loss of *mlaA* or *pldA* – both involved in OM lipid asymmetry – restored growth and morphology in strains lacking *ldtJ*. These effects were independent of OM integrity, implicating lipid asymmetry as the key modulator. These findings align with studies in *Vibrio cholerae*, where OM remodeling genes buffer defects caused by cell wall enzyme loss [49], and support broader models of envelope coordination [9,50].

Transcriptomic profiling revealed widespread gene dysregulation in Δ*ldtJ*, including strong downregulation of *dadA* and *alr* – enzymes essential for D-alanine metabolism and PG biosynthesis [39,40] and upregulation of *argO*, an arginine exporter linked to nitrogen stress [41]. These changes likely disrupt PG precursor synthesis by altering glutamate and *m*DAP availability, contributing to envelope stress. Notably, these transcriptional imbalances were reversed in the Δ*ldtJ* Δ*mla* mutant, suggesting that Mla disruption restores metabolic homeostasis.

Further analysis revealed upregulation of stress-associated genes in Δ*ldtJ*, including *dksA*, and increased ppGpp accumulation, consistent with activation of the stringent response [42,51]. This response, likely triggered by amino acid and PG precursor imbalances, is known to reduce growth and cell size [52,53], mirroring Δ*ldtJ* phenotypes. Restoration of *dksA*, *dadA*, *alr*, and *argO* expression in the double mutant underscores the regulatory link between OM lipid asymmetry and PG homeostasis.

In addition, Δ*ldtJ* cells upregulated a broader set of metabolic and transport-associated genes linked to stress adaptation, nitrogen balance, and PG precursor synthesis (**Table S1**). Some of them included amino acid transporters – *putP* (*A1S_1530*), *mgtA* (*A1S_2070*), catabolism – *aspQ* (*A1S_1466*), *aspC* (*A1S_2508*), biosynthesis – *lysC* (*A1S_1142*), *dat* (*A1S_2454*), *ddc* (*A1S_2453*), and a lysozyme-annotated gene (*A1S_2016*) potentially involved in PG turnover. Many of these genes are associated with nutrient salvage, redox balance, or polyamine metabolism, and are commonly induced under envelope or nutritional stress [54–60]. Strikingly, expression of these genes was downregulated in the Δ*ldtJ* Δ*mla* background, further supporting the role of Mla disruption in alleviating envelope stress and rebalancing cellular metabolism.

Taken together, these findings raise the possibility that the transcriptional signatures observed in Δ*ldtJ* may reflect impaired amino acid uptake or availability, and that disruption of the Mla system – by altering OM lipid asymmetry, potentially affecting selective permeability – facilitates nutrient influx and thereby mitigates the associated metabolic stress. This highlights the tight integration of PG synthesis, amino acid metabolism, and OM homeostasis in *A. baumannii*, with the cell reprogramming gene expression to buffer envelope stress. LdtJ thus functions beyond PG crosslinking, contributing to envelope integrity and cellular homeostasis – a regulatory role not previously described in this organism.

Future studies should explore whether non-enzymatic LdtJ variants (e.g., LdtJ_C390S_ or LdtJ_ΔYkuD_) influence metabolic regulation and whether LdtJ physically interacts with other envelope-associated proteins. Additionally, dissecting the mechanistic basis of the LdtJ-Mla interplay – whether through cross-regulation, envelope biophysics, or shared stress signaling will further illuminate the compensatory dynamics between PG remodeling and OM lipid homeostasis. In conclusion, this study reveals a novel link between LD-transpeptidase-mediated PG remodeling and OM lipid homeostasis in *A. baumannii*. Disruption of *ldtJ* activates envelope and metabolic stress responses, while loss of *mla* restores cellular balance. These findings advance our understanding of envelope coordination and suggest new strategies for targeting interconnected stress pathways in Gram-negative pathogens.

## MATERIALS AND METHODS

### Bacterial strains and growth

All strains and plasmids used in this study are listed in **Table S4**. All *A. baumannii* 17978 strains were initially cultured from −80°C stocks on Luria-Bertani (LB) agar at 37°C. When appropriate, kanamycin was used for selection at a final concentration of 25 μg/ml (Kan^25^).

### Construction of single, double and triple genetic mutants

All *A. baumannii* mutations were generated as previously described [61]. Briefly, the recombinase plasmid pAT03 (expressing REC_Ab_) was expressed in *A. baumannii* ATCC 17978. A linear PCR product containing the kanamycin resistance cassette FLP recombination target (FRT) sites and 125-bp regions of homology to *mlaA, mlaE* and *pldA* was transformed into each strain to generate the respective single deletion mutants: Δ*mlaA,* Δ*mlaE* and Δ*pldA*. To construct double mutants (Δ*ldtJ* Δ*mlaA,* Δ*ldtJ* Δ*mlaE*, Δ*ldtJ* Δ*pldA)* and the triple mutant (Δ*ldtJ* Δ*mlaA* Δ*pldA)*, the pAT03 was first introduced into the appropriate single or double mutant backgrounds. A second linear PCR product targeting *ldtJ*, also containing an FRT-flanked kanamycin cassette with 125-bp homology arms, was then transformed into these strains. Transformants were recovered in LB medium, pelleted by centrifugation, and plated on LB agar supplemented with Kan^25^ for selection. All mutations were verified by PCR and Sanger sequencing.

Following mutant construction, the pMMB67EH::REC_Ab_ plasmid was cured as previously described [61]. The pMMB67EH::FLP recombinase, was then introduced into the cured mutants. Cells were recovered in LB and plated on LB agar containing IPTG to induce FLP expression. Successful excision of the kanamycin cassette was confirmed by PCR.

### Construction of LdtJ_C390S_ and MlaA expression clones

The *ldtJ* coding sequence was amplified from *A. baumannii* ATCC 17978 gDNA. To clone in pMMB67EHKn, the PCR product and plasmid were digested with KpnI and SalI restriction enzymes. The reverse primer was designed to introduce a C390S point mutation, replacing the cysteine at position 390 with serine during amplification. The resulting construct LdtJ_C390S_ was confirmed by Sanger sequencing and transformed into the Δ*ldtJ* strain. This strain was used for Western blotting, complementation, and microscopic studies. For induction of LdtJ_C390S_ expression, culture was grown in Kan^25^ overnight in LB, followed by induction with 2 mM isopropyl-β-d-thiogalactopyranoside (IPTG) the next day.

The *mla* coding sequence was similarly amplified from *A. baumannii* ATCC 17978 gDNA and cloned into pMMB67EHKn using restriction enzymes EcoRI and KpnI. The construct was verified by sequencing and transformed into Δ*ldtJ* Δ*mlaA* double mutant. For overexpression studies, the strain was induced with 0.1 mM IPTG in the presence of Kan^25^.

### Construction of LdtJ domain deletion constructs

Four domain-deletion constructs of *ldtJ* were generated to investigate the functional role of specific regions. The first construct, LdtJ_Δ40-146_ contains the N-terminal 39 amino acids, including the predicted export sequence, while deleting residues 40–146 of the N-terminal domain. The remaining three constructs were designed as follows: 1) LdtJ_Δ147-192_; deletion of residues 147-192, LdtJ_Δ193-282_; deletion of residues 193-282 and LdtJ_ΔYkuD_; deletion of the conserved YkuD domain, retaining only the C-terminal 25 amino acids, which include the α-LdtJ binding region. All constructs were synthesized by Twist Bioscience. The DNA fragments were digested with KpnI and SalI and cloned into the pMMB67EHKn expression vector. Constructs were verified by Sanger sequencing and transformed into Δ*ldtJ* strain. Expression was induced with 2 mM IPTG, and the resulting strains were used for Western Blotting, growth curve analysis, and morphology studies.

### Optical density growth curves

Overnight cultures were back-diluted to an initial OD_600_ 0.05 and distributed into a 24-well plate in triplicate biological replicates. For complementation strains, Kan^25^ and IPTG (2 mM or 0.1 mM, as appropriate) were used. Growth was measured using a BioTek SynergyNeo^2^ microplate reader, which recorded OD_600_ readings hourly at 37°C with continuous orbital shaking. Growth curves were plotted in GraphPad Prism 10 (version 8.4.1). Each experiment was independently replicated three times, and one representative data set was reported.

### CFU growth curve

Overnight cultures were initiated from a single colony and grown at 37°C in LB broth. Cultures were then back-diluted in triplicates to an initial OD_600_ 0.05 and incubated at 37°C with shaking. At designated time points, aliquots were collected, serially diluted, and plated on LB agar. For complementation and overexpression strains, Kan^25^ and IPTG (2 mM or 0.1 mM, as appropriate) were included in the both the liquid and solid media. Plates were incubated overnight at 37°C, and the colony-forming units (CFU) were enumerated. Viability curve was plotted using GraphPad Prism 10 (version 8.4.1). Each experiment was independently replicated three times, and one representative data set was reported.

### SDS/EDTA assay

Overnight cultures were streaked onto standard LB agar plates and LB agar supplemented with 0.012% SDS and 0.185 mM EDTA to assess outer membrane integrity. Plates were incubated overnight at 37°C. The following day, images were captured using a biomolecular imager (Amersham ImageQuant 800 by Cytiva).

### Fluorescent NADA staining

Overnight cultures were back-diluted to an OD_600_ 0.05 and grown at 37°C in LB broth until reaching mid-logarithmic growth phase. Complementation strains were grown in the presence of Kan^25^ and IPTG as required. Cells were harvested, washed once with LB, and resuspended in fresh LB. 2 μL of 10 mM NADA (NBD-(linezolid-7-nitrobenz-2-oxa-1,3-diazol-4-yl)-amino-d-alanine; Thermo Fisher Scientific) was added. Cells were incubated with NADA at 37°C for 20 min with shaking in the dark. Following incubation, cells were washed once and fixed with 1X phosphate-buffered saline (PBS) containing a (1:10) solution of 16% paraformaldehyde.

### Microscopy

Fixed cells were immobilized on 1.5% agarose pads and imaged using an inverted Nikon Eclipse Ti-2 widefield epifluorescence microscope equipped with a Photometrics Prime 95B camera and a Plan Apo 100× 1.45-numerical-aperture lens objective. Green fluorescence images acquired using a filter cube with a 470/40-nm excitation filter and 535/50 emission filter. Image acquisition was performed using NIS Elements software (Nikon).

### Image analysis

All microscopy images were processed with Fiji. Quantitative analysis was performed with the MicrobeJ plugin. Parameters including cell length, width, area, and fluorescence intensity were measured in MicrobeJ. For each strain, at least 100 cells were analyzed per experiment. Data were plotted in GraphPad Prism 10 (version 8.4.1). Each experiment was independently replicated three times, with one representative data set was reported in the quantification, and one representative image was included in the figure.

### Construction of LdtJ overexpression strains for protein purification

The *ldtJ* coding sequence was amplified from *A. baumannii* ATCC 17978 gDNA. The *ldtJ*_C390S_ variant was generated using a reverse primer designed to introduce the C390S point mutation and append a His_8X_ during PCR amplification. Both amplicons were cloned into the NdeI and BamHI restriction sites in pT7-7Kn expression vector. The resulting plasmids, pT7-7Kn::*ldtJ*_His8X_ and pT7-7Kn::*ldtJ*_C390S*-*His8X_, were transformed into chemically competent *E. coli* DH5α cells for propagation and sequence verification via Sanger sequencing. Verified constructs were transformed into chemically competent *E. coli* C2527 (BL21) cells (New England BioLabs, Inc.) for protein expression, purification, and Western blot analysis.

### Purification of recombinant LdtJ

*E. coli* BL21 cells carrying pT7-7Kn::*ldtJ*_His8X_ and pT7-7Kn::*ldtJ*_C390SHis8X_ were cultured in 1 L and 2 L LB broth, respectively, and induced with 1 mM IPTG at 16°C overnight. Cells were harvested by centrifugation, washed in cold 1X PBS, pelleted, and stored at −80°C overnight. Frozen pellets were thawed on ice and resuspended in 20 mL lysis buffer (20 mM Tris, 300 mM NaCl, 10 mM imidazole, pH 8). Cells were lysed by sonication (Fisher Scientific Model 120 Sonic Dismembrator) for 20 sec on and off for a total of 10 cycles at 80% amplitude. Lysates were centrifuged at 15,428 × *g* for 10 min at 4°C. The supernatant was incubated with HisPur Ni-nitrilotriacetic acid (NTA) resin (Thermo Fisher Scientific) prewashed with lysis buffer, on a rotator for 2 h at 4°C. The mixture was loaded onto a 10 mL gravity-flow column (Thermo Fisher Scientific) and washed sequentially with 20 mL lysis buffer containing increasing concentrations of imidazole (0 mM, 15 mM, and 30 mM). Elution was performed with 8 fractions using 500 μL of elution buffer (20 mM Tris, 300 mM NaCl, 250 mM imidazole, pH 8), incubated with the column for 5 min before each gravity elution. Elution fractions containing protein (as determined by SDS-PAGE) were pooled and dialyzed overnight at 4°C using a 20 kDa MW CO 12 mL capacity dialysis cassette (Thermo Fisher Scientific) in dialysis buffer overnight (10 mM Tris, 50 mM KCl, 0.1 mM EDTA, 5% glycerol, pH 8). Protein purity and identity were confirmed by Western blotting using both α-His and α-LdtJ antibodies. Protein concentrations were determined using the Bradford assay.

### Intact mass analysis of LdtJ

Purified wild-type LdtJ protein was separated on a 4-12% SDS-PAGE gel. The gel was stained with Coomassie Brilliant Blue for 10 min and destained overnight with gentle rocking. Protein bands corresponding to LdtJ were excised using a clean scalpel and transferred into 500 μL of dialysis buffer (10 mM Tris, 50 mM KCl, 0.1 mM EDTA and 5% Glycerol, pH 8). Excised gel pieces were submitted to the Proteomics core facility of UT Southwestern Medical Center for intact mass spectrometry analysis.

### Enzymatic activity assays

Activity assays for wild-type (LdtJ) and the catalytic mutant (LdtJ_C390S_) were performed in a final reaction volume of 50 μL containing 20 mM Tris/HCl (pH 7.5), 100 mM NaCl and where indicated, 10 mM D-lysine. Each reaction included 10 μM of either LdtJ or LdtJ_C390S_. To initiate the reaction, 10 μL PG from *E. coli* BW25113Δ6LDT was added. Reactions were incubated overnight at 37°C. The enzymatic reaction was terminated by boiling for 10 min, followed by overnight digestion with cellosyl. After digestion, samples were again boiled for 10 min, then reduced with sodium borohydride and acidified to pH 4.0-4.5. As a control, PG incubated in pH 7.5 buffer without enzyme was processed in parallel. Muropeptide analysis was performed as described previously [62].

### Western blotting

Western blot analysis was done as previously described [63]. Briefly, proteins were transferred to polyvinylidene difluoride (PVDF) membranes (Thermo Fisher Scientific) following SDS-PAGE. Membranes were blocked in 5% non-fat dry milk in Tris-buffered saline (TBS) for 2 h at room temperature. Primary antibodies were used at the following dilutions: α-LdtJ - 1:500, α-His - 1:500, and α-RpoA - 1:1000, followed by HRP −conjugated α-rabbit-IgG for α-LdtJ and α-RpoA and HRP-conjugated α-mouse for α-His at 1:10,000 (Thermo Fisher Scientific). Signal detection was performed using SuperSignal West Pico Plus Chemiluminescent Substrate (Thermo Fisher Scientific).

### RNA sequencing

Transcriptomic analysis was performed as previously described, with modification [64–67]. Total RNA was extracted from *A. baumannii* ATCC 17978 cultures grown in triplicates using the Direct-Zol RNA miniprep kit (Zymo Research), following the manufacturer’s protocol. RNA samples were submitted to the SeqCenter for Illumina RNA sequencing. The resulting reads were aligned to the *A. baumannii* ATCC 17978 reference genome using CLC genomic workbench software (Qiagen). Gene expression levels were quantified as reads per kilobase per million mapped reads (RPKM), and weighted-proportions fold changes were calculated to compare expression across strains. Baggerley’s test on proportions was used to assess differential expression, and *p* values were calculated by two-tailed *t*-test to determine statistical significance.

### Reverse Transcriptase relative fold PCR

RT-relative fold PCR was done as previously reported [65]. Briefly, overnight cultures were back-diluted to OD_600_ 0.05 and grown at 37°C in LB broth until reaching mid-logarithmic phase. Total RNA was extracted using the Direct-zol RNA Miniprep Kit (Zymo Research) according to the manufacturer’s instructions. Genomic DNA contamination was removed using the TURBO DNA-free Kit (Thermo Fisher Scientific). For each strain, 500 ng of DNase-treated RNA was used for cDNA synthesis using random hexamers and the SuperScript III First-Strand Synthesis System (Thermo Fisher Scientific). Quantitative PCR reactions were assembled with 2 µL of cDNA, 0.2 µM gene-specific primers (listed in **Table S5**; designed using Primer3Plus, Andreas Untergasser), and PowerUp SYBR Green Master Mix (Thermo Fisher Scientific). Reactions were run in technical triplicates using the QuantStudio 3 Real-Time PCR System (Applied Biosystems). The housekeeping gene *rpoA* was used as an endogenous control. No-reverse transcriptase controls were included to confirm the absence of genomic DNA contamination. Relative expression levels were calculated using the comparative Ct method, and data were analyzed using DA2 software (Thermo Fisher Scientific).

### Radiolabeled extraction and detection of (p)ppGpp

Radiolabeled pp(G)pp extraction and detection were performed as previously described [68], with minor modifications. Overnight cultures were back-diluted to OD_600_ 0.05 in LB broth supplemented with 5 μCi/mL ^32^P and incubated at 37°C with shaking for 3 h. Cultures were then normalized to lowest OD_600_ value across samples. For each sample, 500 μL of culture was pelleted and resuspended in 100 μL of 2N Formic Acid. Samples were incubated at room temperature for 15 min, and then frozen at −80°C for 15 min. This freeze-thaw cycle was repeated once more. After centrifugation to remove cellular debris, the supernatant was collected. 10 μL of each sample was spotted on a PEI-Cellulose Thin-layer chromatography (TLC) plate and resolved using a 1.5 M potassium dihydrogen phosphate (KH_2_PO_4_), pH 3.4 solvent system. The TLC plate was exposed to a phosphor imaging screen for 48 h, and signals were visualized using a Storm Molecular Imaging scanner. Spot intensities were quantified using ImageJ (Fiji), and data were plotted using GraphPad Prism 10 (version 8.4.1) software.

## ACKNOWLEDGMENTS

The work was supported by funding from the National Institutes of Health (grants R01AI168159 to J.M.B. and W.V. and R35GM143053 to J.M.B.) and the UK BBSRC (BB/W013630/1 to W.V.). We thank Dr. Daniela Vollmer for the preparation of peptidoglycan.

---

**Figure S1.**
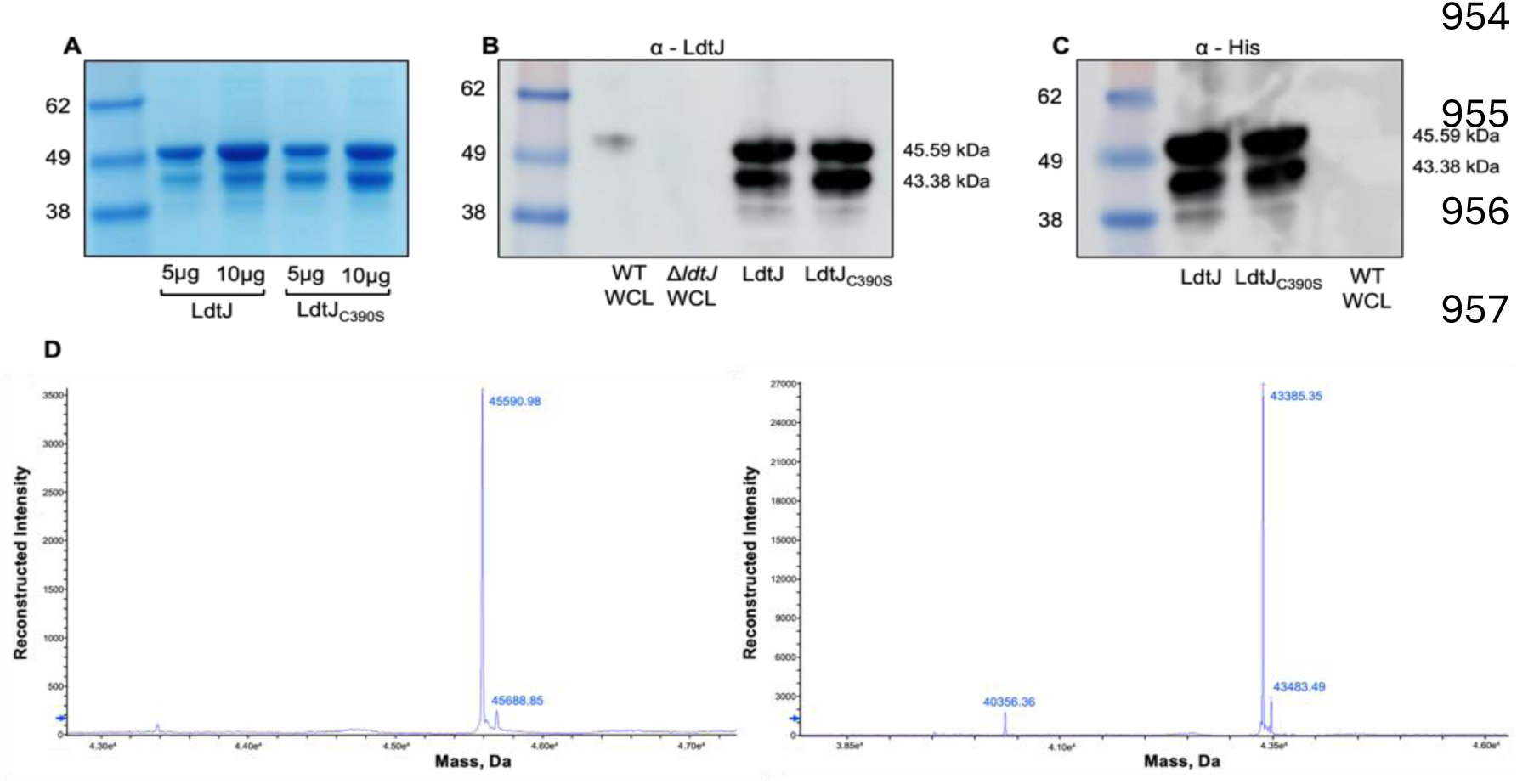
Purification of LdtJ and LdtJ_C390S_. **(A)** Coomassie stained SDS-PAGE gel showing purified recombinant LdtJ and the catalytically inactive mutant LdtJ_C390S_. **(B)** Western blot using α-LdtJ antibody and **(C)** α-His antibody. (WCL = Whole cell lysate) **(D)** Mass spectrometry analysis of purified LdtJ revealed two major products with molecular weights of 45.59 kDa and 43.38 kDa, consistent with the bands observed in panels A–C. Each experiment was independently replicated three times, and representative results are shown.

**Figure S2.**
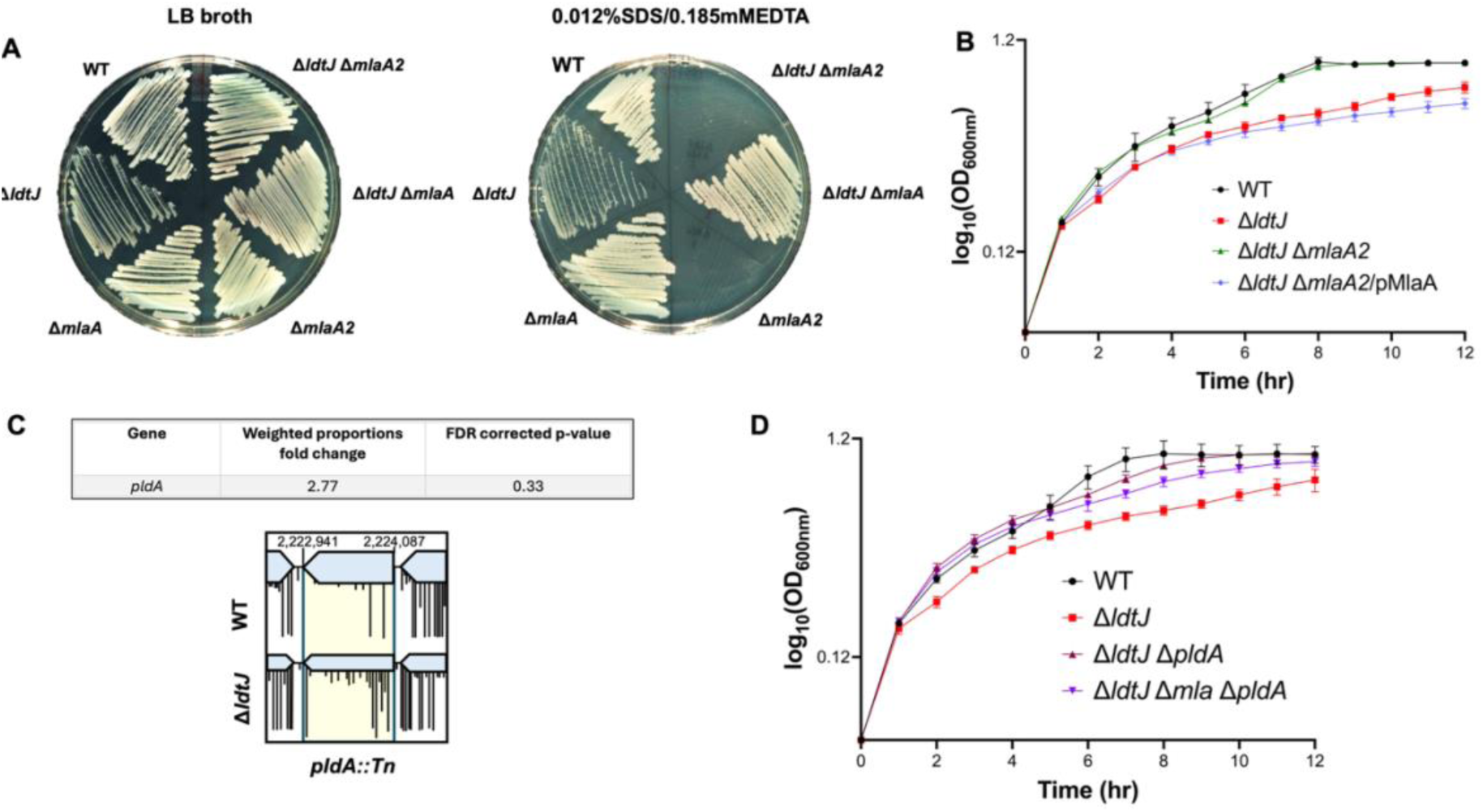
Disruption of outer membrane (OM) asymmetry restores growth fitness in the Δ*ldtJ* mutant. **(A)** SDS/EDTA sensitivity assay of strains with Mla pathway disruption, showing both SDS/EDTA-resistant and −sensitive replicates. **(B)** Growth curve analysis of the SDS/EDTA-sensitive Δ*ldtJ* Δ*mlaA2* strain, demonstrating restored growth fitness like the SDS/EDTA-resistant Δ*ldtJ* Δ*mlaA* strain. **(C)** Transposon sequencing data and insertion profiles for *pldA::Tn* in WT and Δ*ldtJ* backgrounds, indicating significant fold changes in the *pldA* system. **(D)** Growth curve analysis of WT, Δ*ldtJ*, Δ*ldtJ* Δ*pldA*, and Δ*ldtJ* Δ*mla* Δ*pldA* strains. All experiments were independently replicated three times; representative datasets are shown.

## Table Legends

**Table S1: Differential gene expression WT vs. Δ*ldtJ***

**Table S2: Differential gene expression Δ*ldtJ* vs. Δ*ldtJ* Δ*mla***

**Table S3: Differential gene expression WT vs. Δ*ldtJ* Δ*mla***

**Table S4: Strains and plasmids used in this study.**

**Table S5: Primers used in this study.**

## Notes

### Competing Interest Statement

The authors have declared no competing interest.

