## Supplemental figures 1-2 and tables 4 and 5 for "Molecular interplay between peptidoglycan integrity and outer membrane asymmetry in maintaining cell envelope homeostasis"

### Supplementary Figures

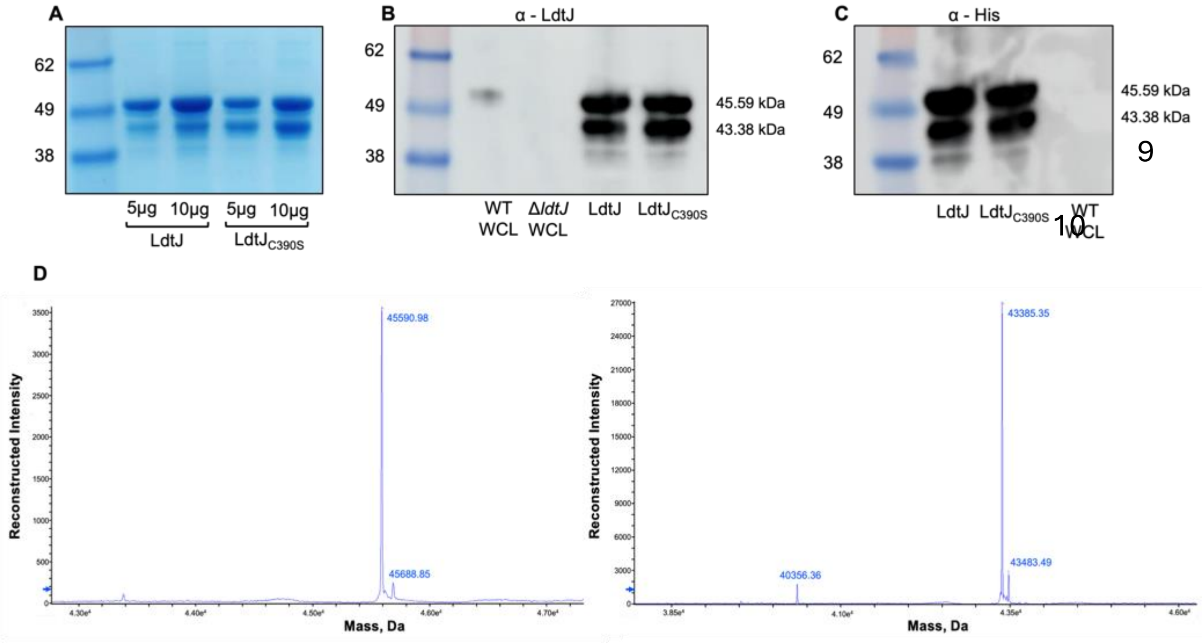


**Figure S1. Purification of LdtJ and LdtJ$_{C390S}$. (A)** Coomassie stained SDS-PAGE gel showing purified recombinant LdtJ and the catalytically inactive mutant LdtJ$_{C390S}$. **(B)** Western blot using α-LdtJ antibody and **(C)** α-His antibody. (WCL = Whole cell lysate) **(D)** Mass spectrometry analysis of purified LdtJ revealed two major products with molecular weights of 45.59 kDa and 43.38 kDa, consistent with the bands observed in panels A–C. Each experiment was independently replicated three times, and representative results are shown.

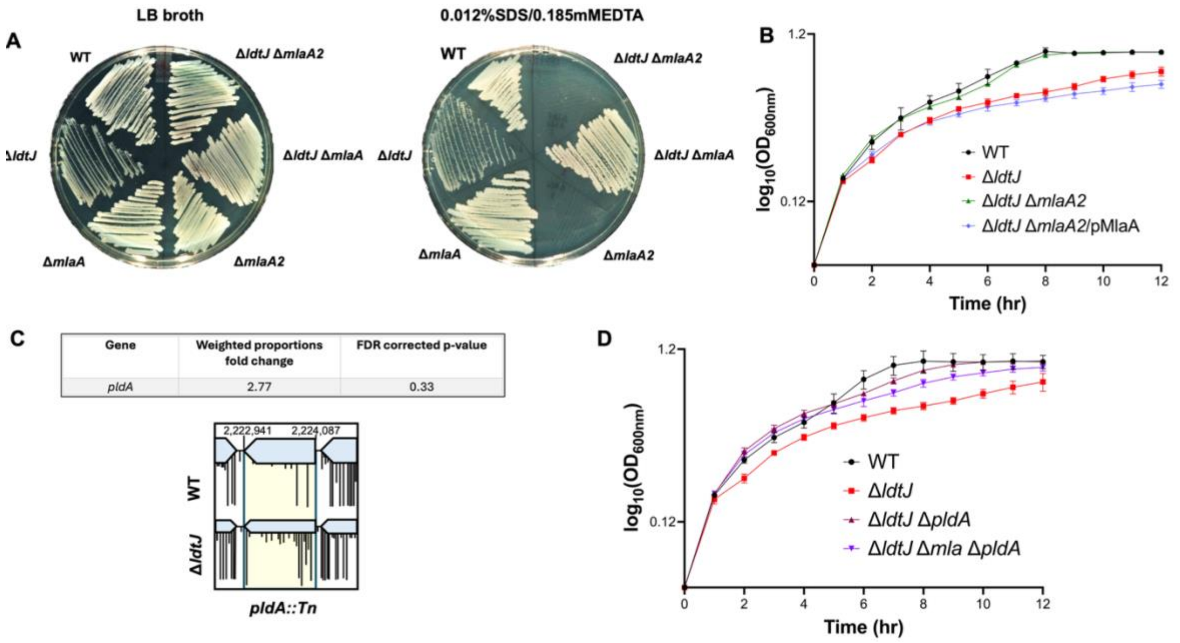


**Figure S2. Disruption of outer membrane (OM) asymmetry restores growth fitness in the Δ*ldtJ* mutant. (A)** SDS/EDTA sensitivity assay of strains with Mla pathway disruption, showing both SDS/EDTA-resistant and -sensitive replicates. **(B)** Growth curve analysis of the SDS/EDTA-sensitive Δ*ldtJ* Δ*mlaA2* strain, demonstrating restored growth fitness like the SDS/EDTA-resistant Δ*ldtJ* Δ*mlaA* strain. **(C)** Transposon sequencing data and insertion profiles for *pldA::Tn* in WT and Δ*ldtJ* backgrounds, indicating significant fold changes in the *pldA* system. **(D)** Growth curve analysis of WT, Δ*ldtJ*, Δ*ldtJ* Δ*pldA*, and Δ*ldtJ* Δ*mla* Δ*pldA* strains. All experiments were independently replicated three times; representative datasets are shown.

**Supplementary Tables**

**Tables S1-3: Please See the Excel Spreadsheet**

**Table S4: Strains and plasmids used in this study.**

| Strain or Plasmid | Description | Reference or Source |
|---|---|---|
| **Strains** | | |
| *A. baumannii* ATCC 17978 | Wild type | [1] |
| *A. baumannii* ATCC 17978 | Δ*ldtJ* (*A1S_2371*) | [2] |
| *A. baumannii* ATCC 17978 | Δ*ldtJ* Δ*mlaA* | This study |
| *A. baumannii* ATCC 17978 | Δ*ldtJ* Δ*mlaE* | This study |
| *A. baumannii* ATCC 17978 | Δ*ldtJ* Δ*pldA* | This study |
| *A. baumannii* ATCC 17978 | Δ*ldtJ* Δ*mla* Δ*pldA* | This study |
| *A. baumannii* ATCC 17978 | Δ*mlaA* (*A1S_0622*) | This study |
| *A. baumannii* ATCC 17978 | Δ*mlaE* (*A1S_3102*) | This study |
| *E. coli* DH5α | Host strain for cloning | [3] |
| **Plasmids** | | |
| pAT03 | pMMB67EH with FLP recombinase, Tet$^R$ | [4] |
| pAT04 | pMMB67EH with Rec$_{AB}$ system, Tet$^R$ | [4] |
| pKD4 | Kan$^R$ | [5] |
| pMMB67EH | pMMB67EH with the Kan$^R$ gene from pKD4 inserted into the PvuI site, Kan$^R$ | [2] |
| pLdtJ | pMMB67EHKn carrying *ldtJ* (*A1S_2371*) | [2] |
| pLdtJ$_{C390S}$ | pMMB67EHKn::*ldtJ*$_{C390S}$ | This study |
| pLdtJ$_{\Delta 40\text{-}146}$ | pMMB67EHKn::*ldtJ*$_{\Delta 40\text{-}146}$ | This study |
| pLdtJ$_{\Delta 147\text{-}192}$ | pMMB67EHKn::*ldtJ*$_{\Delta 147\text{-}192}$ | This study |
| pLdtJ$_{\Delta 193\text{-}282}$ | pMMB67EHKn::*ldtJ*$_{\Delta 193\text{-}282}$ | This study |
| pLdtJ$_{\Delta YkuD}$ | pMMB67EHKn::*ldtJ*$_{\Delta YkuD}$ | This study |
| pMlaA | pMMB67EHKn carrying *mlaA* (*A1S_0622*) | This study |

**Table S5: Primers used in this study.**

| Oligo Name | Sequence (5'-3') |
|---|---|
| **Deletion Primers** | |
| *mlaA* (A1S_0622) Kan-FRT 5' | CAACGATATTTCTATATAATTTGATCAATTTTACATTGATAATA<br>GCTTATCGCTTCTGTATTAAAATATATCTCTACCCAACTGCATA<br>TTGAATATAGCCTGGCTAGCGGTTAAGGAATTATATGAGCGAT<br>TGTGTAGGCTGGAGCTGCTTCG |
| *mlaA* (A1S_0622) Kan-FRT 3' | TTATCAGTGTTATCTTCTGGTACATCTTCAGATTCGTCATCATC<br>AATAAAAGACACATCTGCCGAATCACCTTTTTTCTCGGCAAT<br>CTGGAATGCTTTACGTTGGAGATATAAATCACGAATCATATCC<br>TCCTTAGTTCCTATTCCG |
| *mlaA* confirm 5' | GATCGACTTTACATCTTTGGCACTGC |
| *mlaA* confirm 3' | CTGGTAATGCTTCCAGAGCTTGCT |
| *mlaE* (A1S_3102) Kan-FRT 5' | TCTATCAAATGCATCGTGTAGGGGTAATGTCTTACTCATTATCA<br>CGGTATCAGGTTTATTTATTGGTCTGGTACTCGGATTGCAAGGC<br>TACTCAATATTAGTCAATGTTGGTAGTGAATCAATGAGCGAT<br>TGTGTAGGCTGGAGCTGCTTCG |
| *mlaE* (A1S_3102) Kan-FRT 3' | ACAACCGTGCGGGTCATTGCCGTTGCAATGCCTTCAGGTGTCG<br>GATCACATGCATACCCTTGGAAAACAGCAATCCATGTACACAG<br>CAAAGCAAATACAATGCTCTTAATAATGCCATTTACGACATATCC<br>TCCTTAGTTCCTATTCCG |
| *mlaE* confirm 5' | GCGCTTGATCCCGATTTAATTATGTATGACG |
| *mlaE* confirm 3' | GCGTATAATCAGAACTATAACGTAATTCGTCTAAAGC |
| *pldA* (A1S_1919) Kan-FRT 5' | CAATATTTAATTGAACAAGCACCTGTAAAAGCTGTCGTTGCAC<br>CTTTTGCAAAACGTGATGAATTGCAACAACTGGGTTTTACGAT<br>CAAACAAGTTAATTAAAATAAAAATTGGAGATGAACATGAGC<br>GATTGTGTAGGCTGGAGCTGCTTCG |
| *pldA* (A1S_1919) Kan-FRT 3' | CCTTTAACACTCGCCTATTTAATTGCTACTCCTAAAAAAAGCC<br>GTCTCAAACATCCTCAAGCCAACATTCAAATTGGTTAATAAAA<br>AAGCCGCTGTAAAAGCGGCATTTTTATAGGTCAAACTTAATAT<br>CCTCCTTAGTTCCTATTCCG |
| *pldA* confirm 5' | GGTGGCAAGTATGGACAAACCTG |

| | |
|---|---|
| *pldA* confirm 3' | CCTGTACAGTAGGGCTTTCGCG |
| *ldtJ* (A1S_2371) Kan-FRT 5' | TTATATCCCTTCGCGTCTCAAATAAGCCAATATTAAATTCATAAGAATGAATGATTGGTGAGTTTATGGCCTAAAGGATCTGATTTTCCCTATTGCTTATATGAAAATTCTTAAGGTTGAATTACAGCGATTGTGTAGGCTGGAGCTGCTTCG |
| *ldtJ* (A1S_2371) Kan-FRT 3' | TTAGTAAACCTAGGCTGGTTTTATTTTTATAATCAAAACAATAACTACATATTCCACGGGGCTATGCTAAAAAATTTAATAAAAAAGCCTGCATAAAGCAGGCTCTTTTAATTAAGAGGAATATCCTCCTTAGTTCCTATTCCG |
| *ldtJ* confirm 5' | TACTTGCAGCATGTTACATCGGGTTTA |
| *ldtJ* confirm 3' | GGGTCAGATGCTGAAGCTGAATGGTTA |
| **Complementation Primers** | |
| *mlaA* EcoRI 5' | CGCGAATTCATGAATTATTCTAATTTACTTTTGTCGAGCTTATTAACTGTAGGTC |
| *mlaA* KpnI 3' | CGCGGTACCTTATTTTTCGGTTTTATCAGTGTTATCTTCTGGTACATCTT |
| *ldtJ* KpnI 5' | CGCGGTACCATGTTTGTTCGCTCATTACTCGCTATGAGTTTAAGTTGTATTATTGCTAATGTTGCTTTGGCTGCG |
| *ldtJ*$_{C390S}$ SalI 3' | CGCGTCGACTTATTCTAAGAATTTAACAGTTACGCCTGAACGTACTTTATTACCTAAATCGTTAGCATCCCAGTTCGTTAAACGGATACTACC |
| pMMB67EH confirm 5' | CGGTTCTGGCAAATATTCTGAAA |
| pMMB67EH confirm 3' | CTGCGTTCTGATTTAATCTGTAT |
| **Purification primers** | |
| *ldtJ* NdeI 5' | CGCCATATGTTTGTTCGCTCATTACTCGC |
| *ldtJ* His 8X BamHI 3' | CGCGGATCCTTAATGGTGATGGTGATGGTGATGGTGTTCTAAGAATTTAACAGTTA |

| | |
|---|---|
| $ldtJ_{C390S}$ His 8X BamHI 3’ | CGCGGATCCTTAATGGTGATGGTGATGGTGATGGTGTTCTAAGAATTTAACAGTTACGCCTGAACGTACTTTATTACCTAAATCGTTAGCATCCCAGTTCGTTAAACGGATACTACCGTG |
| pT7-7 confirm 5’ | CGATTCGAACTTCTGATA |
| pT7-7 confirm 3’ | ATCGATGATAAGCTT |
| **Relative fold PCR primers** | |
| *A1S_0095* 5’ | CGGGTATTCACTACGAAAACCG |
| *A1S_0095* 3’ | TGAACCGCTTCCATTTGTGC |
| *A1S_0096* 5’ | AGCGCAAATTGAGTGGGTAG |
| *A1S_0096* 3’ | TCCCAAGCCTGCACATAATG |
| *A1S_1046* 5’ | TGCCGCGTTTAAAACTACCC |
| *A1S_1046* 3’ | CGCAACCGAACCAATCAGTAC |
| *rpoA* 5’ | AATGCGCGTGTAGAACAACG |
| *rpoA* 3’ | CAAGATTGTTGCCGCTTTGC |
